# Altered Interpersonal Neural Synchronization during Social Interaction After Shared Excluded Experiences in Depressed Adolescents

**DOI:** 10.1101/2025.03.24.644779

**Authors:** Yuwen He, Jieting Chen, Yong Lin, Natalia Chan, Fei Gao, Lulu Liu, Xiaoqing Yin, Yao Sun, Minghui Li, Sifan Zhang, Zihan Wei, Liangxuan Yu, Xinyi Huang, Zhihai Su, Zhen Yuan

## Abstract

**Background:** Major depressive disorder (MDD) is common in adolescents, and the special development stage, during which adolescents’ brain and neuroendocrine system develop intensively, makes it subtly difficult to develop prevention and treatment strategies for depressed adolescents compared with depressed adults. Meanwhile, public psychosocial stressors significantly influence adolescents’ mental health and social interaction, rendering it essential to explore how a shared psychosocial stressor, i.e., shared excluded experiences, influences social interaction in depressed adolescents.

**Methods:** We designed a 4-player cyberball game to probe adolescents’ responses to shared excluded experiences and explore the underlying interpersonal neural synchronization (INS) with functional near-infrared spectroscopy (fNIRS).

**Results:** We found that shared excluded experiences could enhance adolescents’ social interaction preferences but decreased INS in each pair of excluded adolescents, which indicates a reduced willingness to interact with others after the exclusion. However, no significantly different behavioral responses to the shared excluded experiences were found in depressed adolescents compared to adolescents as healthy controls (HC). Further analyses revealed that adolescents with MDD experienced more negative feelings than HC after exclusion. Of note, adolescents with MDD demonstrated stronger INS than HC, indicating the potential empathic stress in depressed adolescents. In addition, there existed altered brain-behavioral association patterns in responses to shared excluded experiences in depressed adolescents.

**Conclusions:** In summary, our study gives us deeper insights into how a shared psychosocial stressor impacts the INS in depressed adolescents, and it might be demonstrated that INS could be more sensitive than behavioral responses to detect social interaction deficits in depressed adolescents.

## Introduction

Major depressive disorder (MDD) is common among adolescents, bringing high suicide risk among adolescents and great burdens to families and societies (1). In addition, the prevention and treatment strategies for depressed adolescents are slightly different from those for depressed adults. It warrants more intensive efforts to understand the potential deficits in depressed adolescents, facilitating efficient prevention and treatment approaches to improve the mental health environment for adolescents. Many risk factors contribute to the high rate of depressive disorders in adolescents, including genetic risk, family history of depression, progressively developing periods of brain prefrontal cortex-related circuits and hormonal system, and psychosocial stressors (1–3). Among them, exploring how psychosocial stressors play out on the prefrontal cortex-related circuits can help us develop prevention and treatment approaches for depressed adolescents in different social communities (i.e., clinical service, educational system, and residential community).

Public psychosocial stressors, i.e., the COVID-19 pandemic, terrorist attacks, gun shootings, or car attack accidents, would have a tremendous impact on the individuals (4, 5). These public stressors would increase the vulnerability of getting depressed or relapsing into depressed status (6). However, how these psychosocial stressors shared by people affect individuals’ cognitive and affective functions, especially in depressed adolescents, was scarcely examined. Social deficits are common in individuals with major depressive disorder (7). Exploring the relationship between psychosocial stressors and social functions would help us to understand what kinds of declines might relate to the occurrence of depression. Surprisingly, it is little known whether the shared negative experiences would expose the social function of depressed individuals more vulnerable. Thus, it would be necessary to design a laboratory task generating shared psychosocial stressors to explore how it would influence social functions in depressed individuals. Of note, social rejection or social exclusion is a common risk factor for depression (1), and it can be easily manipulated via laboratory tasks (8). Meanwhile, social preference during interpersonal interaction is important in forming social relationships (9). So, we designed a multiplayer cyberball game to generate social exclusion for two participants simultaneously and examined how this shared psychosocial stressor would influence interpersonal interaction preferences in depressed individuals.

Depressed individuals are more sensitive to social exclusion (10). Previous studies suggested individuals might have different responses to social exclusion. Some people might seek re-acceptance by the excluder, while others may reduce interaction with the excluders (11, 12). Meanwhile, some studies reported that shared negative emotions may facilitate interpersonal relationships (13), which indicates that individuals prefer to interact with the ones sharing negative experiences with them.

This conflicting evidence suggests potentially diverse individual differences in social interaction preferences. Social withdrawal is common in individuals with MDD (7). However, it is not known whether the social interaction preference is altered in depressed individuals, given their low interaction interests, compared to healthy controls (HC). The potentially diverse individual interaction preferences may give us a better chance to examine the potential differences in interaction preferences between depressed adolescents and the HC.

Simultaneous recording of brain signals in several individuals enables us to probe the underlying interpersonal neural synchronization (INS) of interpersonal interaction (14, 15). Such biological underpinnings of social interaction might be more stable phenotypes to assess the social deficits in depressed individuals. However, there appears no study examined the potentially altered INS in MDD compared with HC with fNIRS or electroencephalography (EEG) techniques, except one research reported the association between INS and depressive symptoms (16). This might relate to the fact that cognitive and affective deficits are not the primary interventional targets in the conventional medicine system (17). However, the improvement of cognitive and affective deficits are significant predictors of the well-being of depressed individuals (18). Thus, more efforts are warranted to develop efficient approaches to assess the social deficits in depressed individuals, which would be important for developing better interventional methods to enhance the well-being of depressed individuals. In the current study, we explored how a shared psychosocial stressor, social exclusion, might modulate interpersonal neural interaction during a social task performance in depressed individuals.

In summary, our study aimed to 1) adopt, and modify a paradigm that can generate a shared psychosocial stressor, i.e., social exclusion, and examine how this shared psychosocial stressor would influence the social interaction preference in depressed adolescents; 2) how this shared psychosocial stressor modulates the INS during the social task performance in depressed adolescents; 3) whether there are altered social interaction preferences in depressed adolescents compared with HC; 4) whether there are altered INS during this social task performance in depressed adolescents compared with HC.

## Methods and Materials

### Participants

Thirty-six pairs of right-handed healthy adolescents were recruited from the local community of Macau and two middle schools in Zhuhai, China. All the healthy adolescents did not report any history of diagnosis of any mental disorders. Forty pairs of right-handed adolescents with MDD were recruited to the study in the Department of Psychiatry, Fifth Affiliated Hospital, Sun Yat-sen University. All individuals with MDD had met the diagnostic criteria designated by the Diagnostic and Statistical Manual of Mental Disorders, 5th Edition (DSM-5) and were diagnosed by professional psychiatrists. Several pairs were excluded due to incomplete data collection (refer to supplementary materials). Finally, 34 pairs of adolescents with MDD and 34 pairs of matched controls were included for further analysis.

### Paradigm

The task setting is shown in Fig 1. A. Two participants (both of them were either individuals with MDD or healthy controls) were required to play a modified four-player Cyberball game (Fig 1. C & D) wearing an fNIRS headband, while the other two were virtual players programmed by the computer. The participants were asked to pass the ball to one of the other players once they received it as they wished. There were three stages, including the acceptance stage, exclusion stage, and re-acceptance stage, which was further divided into three sessions.

**Fig 1.**
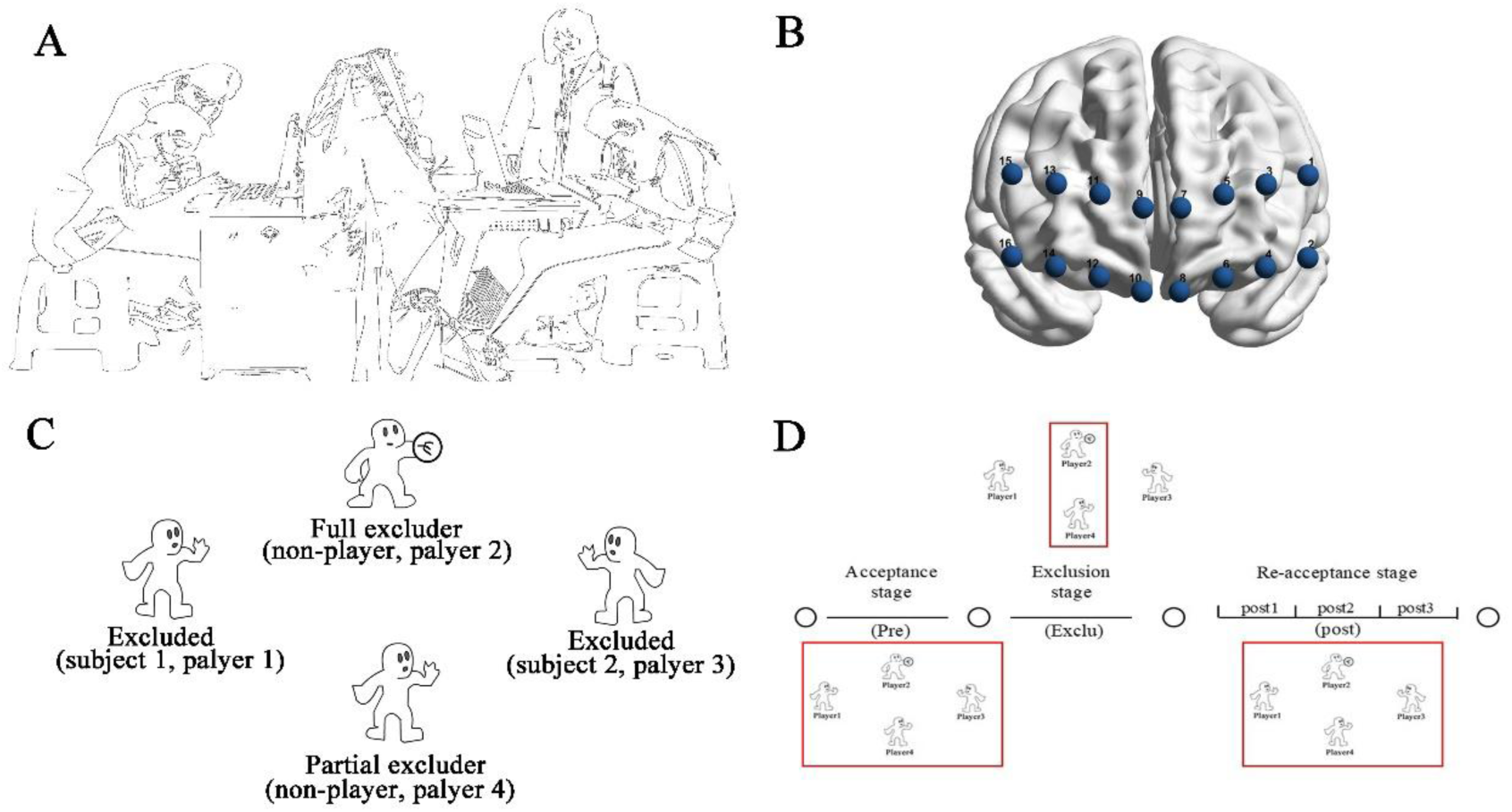
Behavioral paradigm setups. A: behavioral setups for the experiment; B: the distribution of optical channels; C: the modified 4-player Cyberball game; D: the procedures of the paradigm. The red rectangle indicates engaging players on a particular stage.

### Behavioral measurements

Hamilton depression scale (19) was administered to each participant in a structured interview conducted by trained experimenters, assessing the severity of depressive symptoms experienced in the past week. A digital span test (20) was also performed for each participant to measure their verbal working memory. In addition, the participants were asked to fill out a need threat scale (21) and a developed positive attitude scale for all the other players (detailed in supplementary materials) to assess the psychological harm and impact on the interpersonal relationship.

### Behavioral statistics of the modified four-player Cyberball game

Several behavioral statistics were used to characterize the task performance of each pair of participants (detailed in supplementary materials). Two statistics were used to assess social interaction preferences, including the mean pass rate and reciprocal pass rate. The differential pass rate and the differential reciprocal pass rate were used to quantify the differential strategy used by the two players of a dyad. The pass rate skewness and mean switch rate were used to assess the overall degree of biased selection preference.

### Interpersonal neural synchronization

The fNIRS data acquisition and preprocessing procedures were detailed in the supplementary materials. Wavelet coherence was used to index the INS (22). Calculating the wavelet coherence for each pair of time series can generate coherence degrees for each time point in each frequency. The INS for each pair of channels between two participants was calculated before the INS was transformed into the Fisher-Z score. It was followed by averaging the INS across time points for each frequency. There were 16*16 pairs of channels between participants. Since the participants in a dyad were randomly matched, it was assumed there were no different roles between the two participants. Thus, the INS in the upper and the lower triangular matrix of channel pairs would be averaged.

To reveal the frequency range and channels related to the social exclusion experience, we compared the Fisher-Z score of INS in each frequency in each channel pair between the exclusion stage and acceptance stage or between the re-acceptance stage and acceptance stage with a t-test. A permutation method with a centered threshold *p* = 0.001 and a neighboring threshold *p* = 0.05 to select the channels and frequency range related to exclusion experience (23). Finally, 31 pairs of channels were selected, and the frequency range between 0.01-0.019 Hz was revealed. The Fisher-Z score of INS in the abovementioned selected frequency range was averaged across time and frequency for each stage in each pair of channels in each pair of participants.

### Statistical analysis for the behavioral responses to the shared excluded experiences

The responses to shared excluded experiences were first examined in the HC group. The statistics, including reaction time, mean pass rate, mean reciprocal pass rate, differential pass rate, differential pass rate, pass rate skewness, and mean switch rate, were compared between the acceptance and re-acceptance stages with the paired-sample t-test.

### Statistical analysis for the interpersonal neural responses to the shared excluded experiences

The manipulative effects of INS in the selected pairs of channels were examined with the repeated-measurement ANOVA across the HC and MDD groups. The significant effects should survive false discovery rate correction (FDR q < 0.05). In addition, the post hoc analyses used paired-sample t-tests to examine the manipulative effects of shared excluded experiences in INS in HC and MDD groups separately.

Correlation analysis was also performed to examine whether the changed INS in channels showing significant manipulative effects of shared excluded experiences correlated with the altered behavioral responses between the sub-session of the re-acceptance stage and the acceptance stage.

### Statistical analysis for between-group differences in behavioral responses

The behavioral responses to shared excluded experiences were examined in the MDD group with paired sample t-tests before performing between-group analysis. Two-sample t-tests were used to examine the between-group differences in the abovementioned behavioral statistics in the acceptance and re-acceptance stages. Two sample t-tests were also used to examine the between-group differences in behavioral measurements including the need threat scale and positive attitude scale for each player. Additionally, linear mixed models were used to examine whether the severity of depression correlates with the behavioral measurements.

### Statistical analysis for between-group differences in interpersonal neural synchronization to shared excluded experiences

The INS in the selected pairs of channels in each session of the re-acceptance stage was compared between HC and MDD groups with the two-sample t-test. In addition, the group (HC & MDD) * stage (re-acceptance & acceptance) interaction effect was examined with repeated-measurement ANOVA. Significant between-group effects should survive FDR correction (*q* > 0.05).

The channels showing between-group effects in the re-acceptance stage were also examined to reveal whether the INS in the corresponding session was associated with behavioral measurements with correlation analysis in the HC and MDD groups separately. Correlation analysis was also performed to examine whether the change in INS, in the channels showing significant group*stage interaction effects, was associated with the change in behavioral responses between acceptance and re-acceptance stages in two groups separately. In addition, a linear regression model shown below was conducted to examine whether the diagnostic groups modulated the association between INS and behavior.

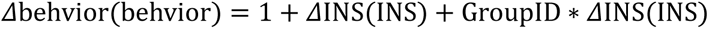

Dependent variable (Δbehvior(behvior)): the change in behavior or the behavior after exclusion; ΔINS(INS): the change in INS between re-acceptance and acceptance stages or the INS in the re-acceptance stage.

## Results

### Demographic data

The average age of healthy adolescents was 17.87 years, while that of depressed adolescents was 17.21 years (Table 1), showing no between-group differences (*p* = 0.11). The HC and MDD groups were also matched in sex (*p* = 0.09), education (*p* = 0.07), and digital working memory (*p* = 0.12).

**Table 1.**
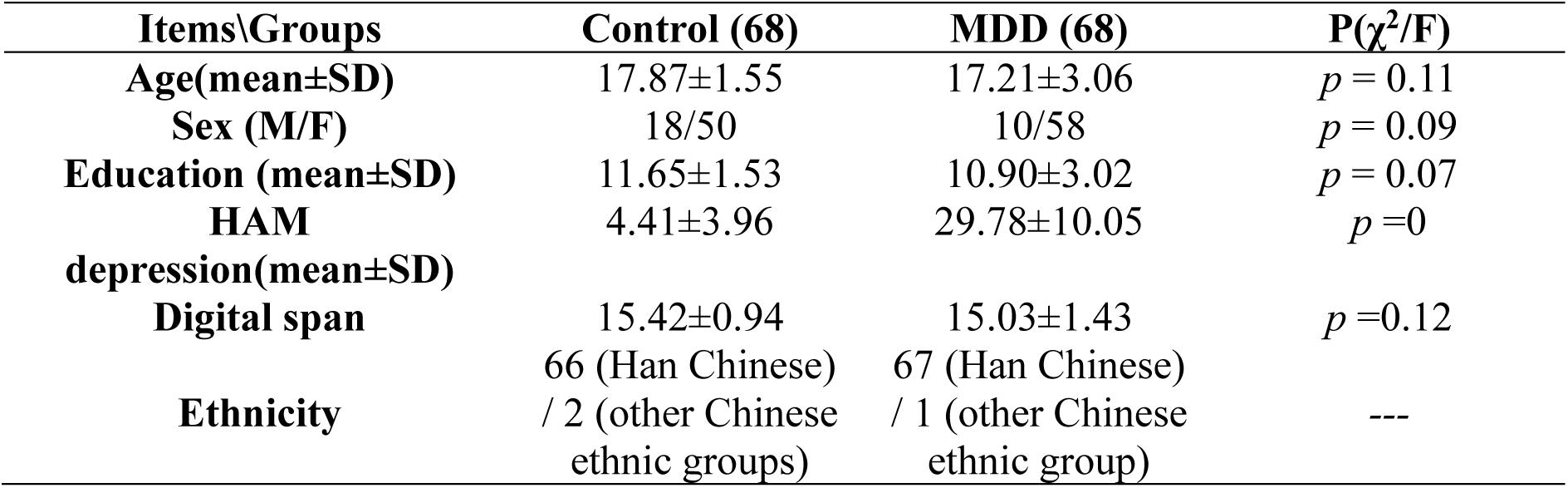
Demographic data and behavioral assessments.

### Behavioral response patterns to shared excluded experiences in healthy adolescents

The HC and MDD groups were also matched in age, sex (*p* = 0.09), education (*p* = 0.07), and digital working memory (*p* = 0.12). Healthy adolescents increased interaction preference to the fellow participant sharing the excluded experience indicated by significantly increased pass rate (*p* < 0.01) and reciprocal pass rate (*p* < 0.01) to the other excluded participant (Fig 2. A & B) after the shared excluded experiences, while reduced interaction preference to the excluders, as indicated by reduced pass rate to the excluders (Fig 2. A, partial excluder: *p* = 0.14; full excluder: *p* < 0.001). Meanwhile, healthy adolescents revealed reduced strategy difference to partial excluder (*p* = 0.03), reduced strategy difference of reciprocal pass to the excluded fellow participant (*p* = 0.04) after shared social excluded experiences, while no significant reduction of strategy difference to excluded fellow participants and full excluder in overall pass rate (Fig 2. C & D). In addition, it was found that shared excluded experiences enhanced the biased selection preference in healthy adolescents as indexed via the pass rate skewness (Fig 2. E) and switch rate (Fig 2. F). Of note, these results also indicated that shared excluded experiences did not cause the excluded participants to form another exclusive circle when they were able to do so.

**Fig 2.**
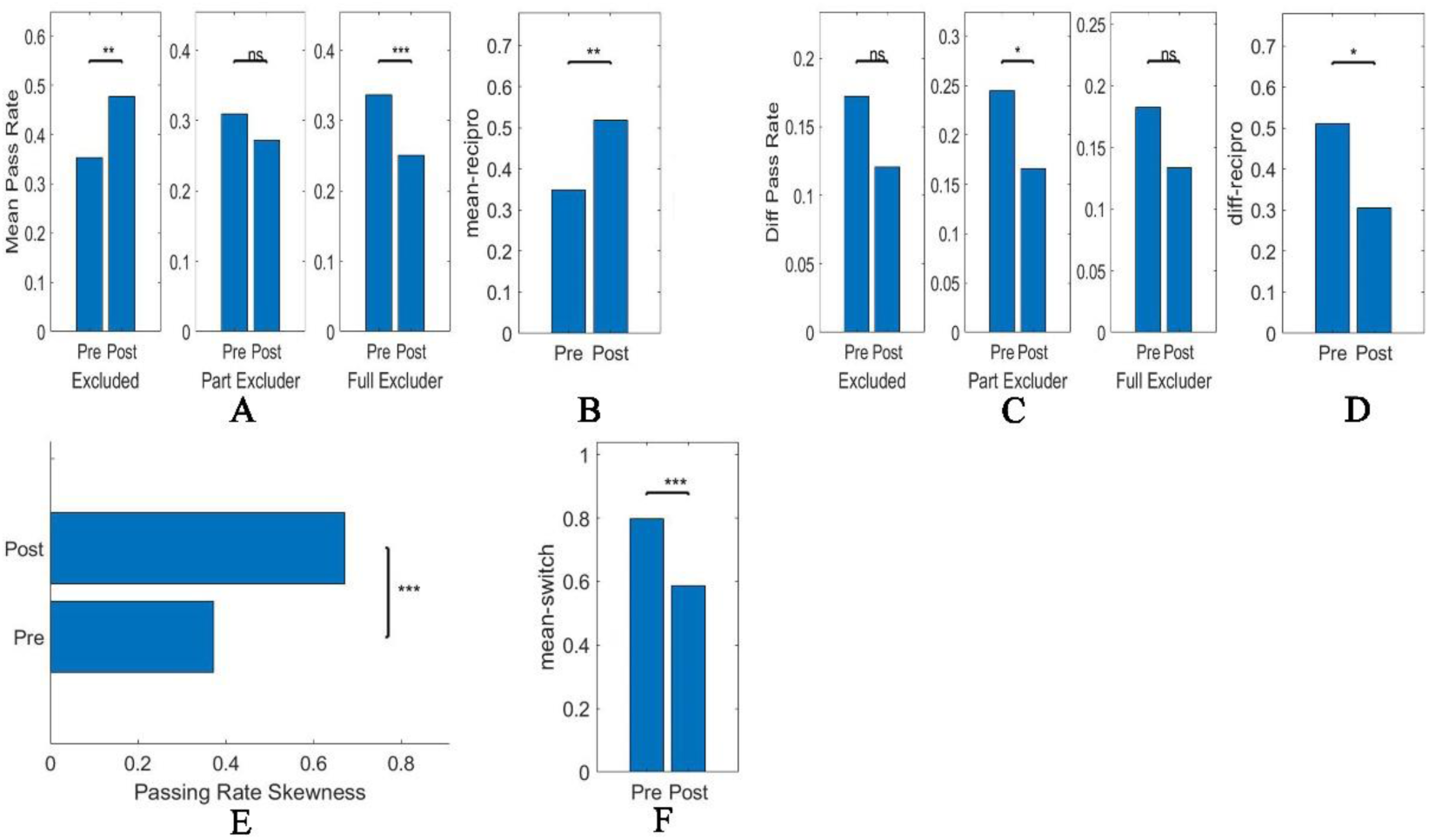
Behavioral responses to shared excluded experiences in healthy adolescents. A-B: the social interaction preference indexed by the mean pass rate and reciprocal pass rate significantly changed after shared excluded experiences; C-D: the strategy difference indexed by the differential pass rate and differential reciprocal pass rate significantly changed after shared excluded experiences; E-F: the selection preference indexed by the pass rate skewness and switch rate significantly changed after shared excluded experiences. Denotes: pre: before social exclusion; post: after shared excluded experiences; Excluded: another excluded player; Part Excluder: partial excluder; mean-recipro: mean reciprocal pass rate; diff pass rate: differential pass rate; diff-recipro: differential reciprocal pass rate; mean-switch: switch rate. *: *p* < 0.05; **: *p* <0.01; ***: *p* < 0.001

### Interpersonal neural synchronization modulated by shared excluded experiences

The procedures for calculating INS are shown in Fig 3. A. Five interpersonal channels in the 2nd session and two interpersonal channels in the 3rd session of the re-acceptance stage displayed significantly lower INS when compared with the acceptance stage (Fig 3. B & C-I, FDR *q* < 0.05), suggesting the social exclusion experience might reduce the adolescents’ willingness to interact with others. Detailed comparisons within the HC or MDD groups are shown in Fig supp 1, which indicates healthy and depressed adolescents revealed similar interpersonal neural responses after shared excluded experiences. The involved interpersonal channels include that from one’s right middle frontal gyrus (ch15, MFG) to the other’s left superior frontal gyrus (ch5, SFG), left MFG (ch3), right orbital frontal gyrus (ch10, OFG), from one’s right OFG (ch10) to the other’s left MFG (ch3), and from one’s left OFG to the other’s left SFG (ch5).

**Fig 3.**
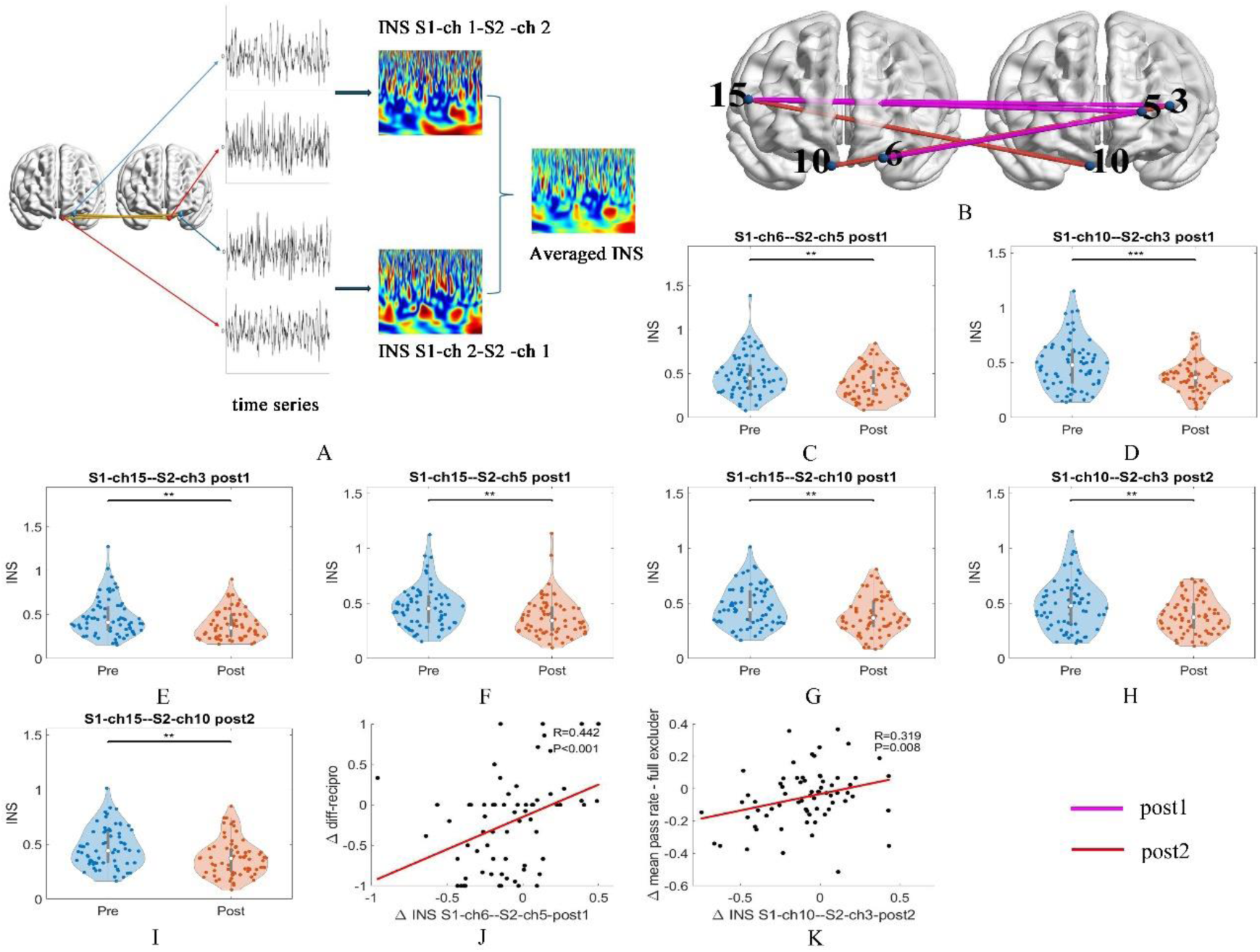
The interpersonal neural responses and their associations with behavioral responses after the shared excluded experiences. A: the procedures to calculate INS; B: seven INSs impacted by shared excluded experiences; C-I: the violin plots of comparisons of INS between the acceptance stage (pre-social exclusion) and the 1st or 2nd session of the re-acceptance stage (post-social exclusion) across HC and MDD; J-K: the association between change in INS and change in behavioral responses across groups.

The INS from one’s left OFG to the other’s left SFG significantly correlated with the change in differential reciprocal pass rate in the 1st sub-session of the re-acceptance stage (Fig 3. J, FDR corrected *q* < 0.05), indicating this interpersonal neural disconnection underlined the reduced differential reciprocal pass rate between participants after social exclusion. Meanwhile, the INS from one’s right OFG to the other’s left MFG significantly correlated with the changed pass rate to the full excluder (Fig 3. K, FDR corrected *q* < 0.05), indicating this interpersonal disconnection underlined the reduced pass rate to the full excluder.

### Between-group differences in behavioral responses and measurements to the shared excluded experiences

The depressed adolescents revealed similar behavioral responses to shared excluded experiences as did the healthy adolescents (Fig 4). As shown in Fig 4. A, the reaction time in the MDD group was higher than that in the HC group before (*p* < 0.01) and after social exclusion (*p* < 0.001). However, no other between-group effects were shown in any behavioral responses at any stage (Fig 4. B-G).

**Fig 4.**
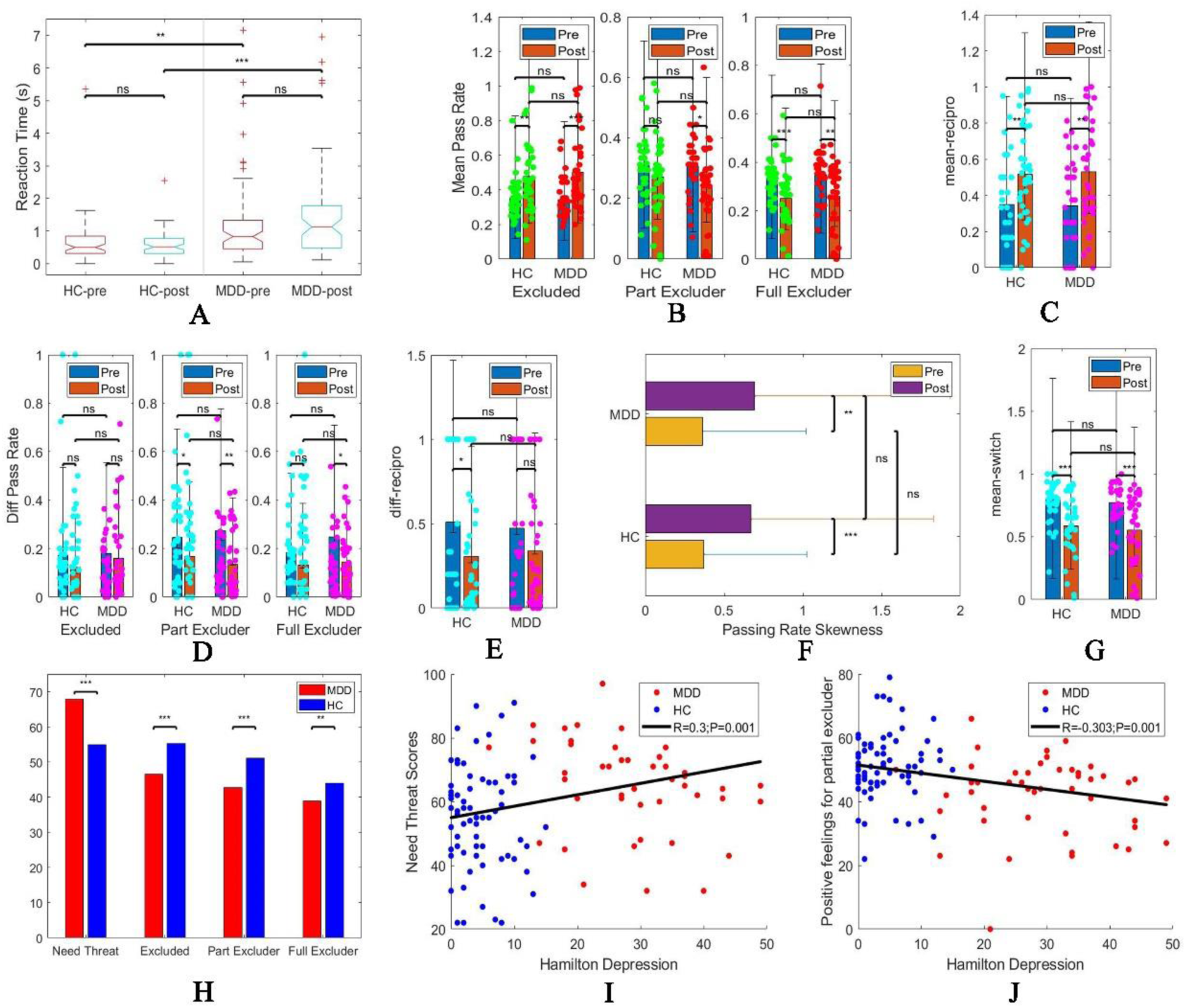
Between-group effects of behavioral measurements. A-G: the within-group (manipulative) and between-group effects in reaction time, social interaction preference (indexed by mean pass rate, reciprocal pass rate), strategy difference (indexed by the differential pass rate, differential reciprocal pass rate), and selection preference (indexed by the pass rate skewness and switch rate). H: between-group comparisons of need threat, and the degree of positive feelings towards other players; I: Association between Hamilton depression and the need threat score; J: Association between Hamilton depression and the degree of positive feelings towards the partial excluder.

Compared with HC, adolescents with MDD had higher scores on the need threat scale (Fig 4. H), indicating they were hurt more than HC after the exclusion experience. In addition, depressed adolescents had less positive feelings towards all the other players (Fig 4. H). The severity of depression significantly correlated with the need threat score and the positive feelings for the partial excluder (Fig 4. I & J).

The time or group*time interaction effects of mean pass rate among the three sub-sessions of the re-acceptance stage (Fig supp 2) indicated there were potential behavioral fluctuations after social exclusion in adolescents.

### Between-group differences in interpersonal neural synchronization

The INS from one’s right MFG (ch13) to the other’s left MFG (ch3) was significantly higher in individuals with MDD compared to HC in the third session of the re-acceptance stage (Fig 5. A & B), while this difference was not observed before the social exclusion experience. Further analyses revealed that higher INS in the HC group correlated with more positive feelings to the other excluded one (Fig 5. C, R=0.409, p =0.018), while there appeared a trend of negative correlation in the MDD group (Fig 5. C, R=-0.203, p =0.242). Further linear regression analysis revealed a significant modulating effect by groups to these associations (β = -10.54, *p* < 0.001). These results might suggest the higher INS in depressed adolescents compared to healthy adolescents might reveal their struggle to show friendly interaction with others, but they had reduced willingness to interact with others. In addition, a significant correlation was observed between the change in INS and the change in the differential pass rate to the partial excluder in the MDD group (Fig 5. D, R=0.362, p =0.036). In contrast, this pattern was not observed in the HC group (Fig 5. D, R=0.059, p =0.738). However, further linear regression analysis did not reveal a significant interaction effect (β = 0.19, *p* = 0.21).

**Fig 5.**
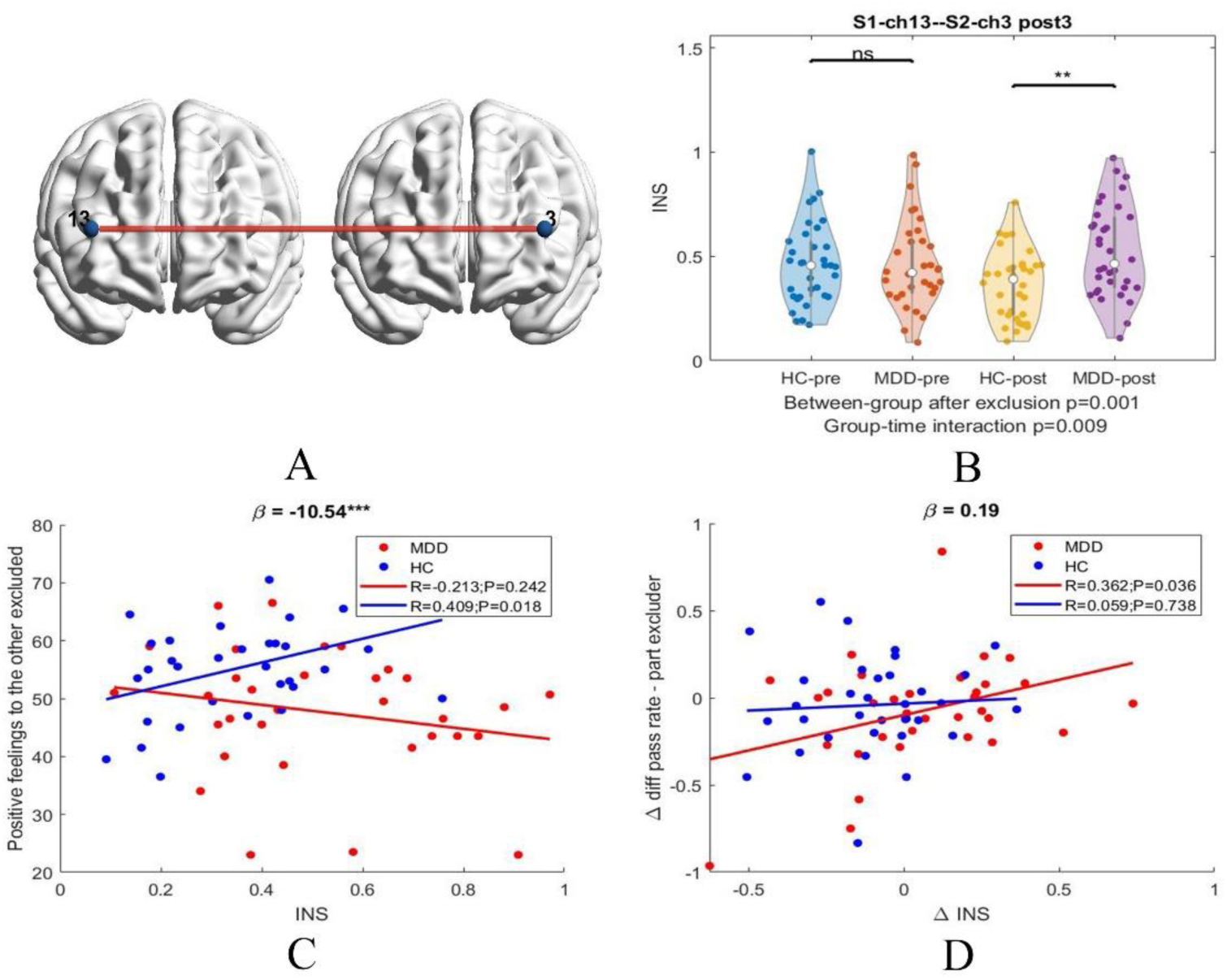
Between-group comparisons of INS and the associations, modulated by diagnostic groups, between INS and behavioral measurements. A-B: INS from one excluded participant’s right MFG (ch 13) to the other excluded participant’s left MFG (ch 3) showed a significant between-group effect; C: the association between INS and the score of positive feelings to the other excluded participant was modulated by diagnostic groups (*β* = -10.54, *p* < 0.001); D: the different associations between change in INS and change in the differential pass rate to the partial excluder in the HC and MDD groups (*β* = 0.19, *p* = 0.21).

## Discussion

Our study examined how the shared psychosocial stressor, namely the shared excluded experiences, influences social interaction preferences between depressed or healthy adolescents. It was demonstrated the shared excluded experiences facilitate the interaction preference between the excluded adolescents in the HC and MDD groups, but they do not necessarily lead to the formation of exclusive subgroups that hinder larger group interactions. The INS between adolescents who shared excluded experiences was significantly reduced, indicating the excluded experiences negatively affected the adolescents by significantly diminishing their willingness to engage with others, leading to a decrease in the INS meantime. These reduced INSs corresponded with the reduced interaction with the excluder and the diminished differential strategies among the excluded adolescents. Compared with HC, the adolescents with MDD did not reveal any altered patterns of behavioral responses. However, they tended to experience more negative feelings from the social exclusion experience and less positive feelings towards others. In addition, the adolescents with MDD displayed higher INS between one’s right MFG and the other’s left MFG than HC. Further analysis revealed that diagnostic groups modulate the associations between INS and behavioral assessments, which would help differentiate adolescents in the HC or MDD group.

Our study found that the shared excluded experiences can facilitate social interaction preferences and reduce the strategy differences between the excluded adolescents in both the HC and MDD groups. In line with previous studies observing that sharing negative experiences would enhance emotional empathy and social bonding (24), shared excluded experiences might induce mutual empathy and facilitate interpersonal bonding. As revealed by a previous study, sharing distinctive negative emotions had different beneficial effects on individuals (13), indicating that shared exclusion experiences might impact social bonding in unique ways. Specifically, forming a close bond with the other excluded ones might give them another chance to obtain affiliation (25), while another study suggested that sharing social exclusion experiences can facilitate bonding stemming from the perceived similarity (26).

However, it is arguable that the shared exclusion experiences would not encourage the excluded individuals to form another exclusive circle, which might be similar to the finding that past adversity predicts increased empathy and compassion for others (27).

In contrast with the behavioral responses, the INS between the excluded adolescents was reduced after the social exclusion, suggesting the excluded adolescents might significantly reduce their willingness to interact with others after the exclusion experience even though the excluded individuals prefer to interact with the other excluded ones than the excluders. Our results are consistent with several previous studies that social exclusion experiences enhance an individual’s social withdrawal tendency (28, 29), which might be the underlying reduced INS between excluded adolescents. Meanwhile, these results correspond to the finding in a previous study that shared exclusion experiences facilitate social bonds less strongly than shared acceptance experiences (26). In other words, despite the shared exclusion experiences can promote bonding, which would be marred by diminished willingness to interact with others triggered by exclusion experience. In our study, the widespread reduced INS among the excluded adolescents might argue that the drive to commit social withdrawal outweighed the developed preference to interact with the other excluded ones. However, our study cannot provide more straightforward evidence on this aspect due to the force selection feature in this paradigm. The correlation results indicate that reduced INS correlates with reduced strategy difference between the excluded ones, which is similar to the findings suggesting sometimes negative emotions have the power to facilitate performance (30, 31). It was also observed that reduced INS correlates with a reduced pass rate to the excluder, which might complement the results observed in a previous study that the more upset after viewing a social rejection event, the more likely they would punish the excluder (32).

There were no significant between-group differences in behavioral responses to the shared excluded experiences, which was out of our expectations. We assumed depressed adolescents might have relatively intact behavioral responses to shared excluded experiences. However, the adolescents with MDD had higher INS between one’s right MFG and the other excluded one’s left MFG compared to healthy adolescents after shared excluded experiences. Previous studies demonstrated that individuals with MDD had higher empathic stress (33), which might be the reason they ostensibly care more about the excluded fellow participant and have higher INS when they have a reduced willingness to interact with others. Thus, more effort is necessary to help depressed adolescents deal with psychosocial stressors than healthy adolescents since there might be higher empathic stress in depressed adolescents. The MFG is a vital brain region engaged in executive function and decision-making (34), which might signify the altered neural interaction related to decision-making during social interaction in adolescents with MDD. Meanwhile, our study revealed the altered INS but no significantly different behavioral responses to shared excluded experiences in depressed adolescents, which might suggest interpersonal neural signals are more sensitive to detecting the potential social interaction deficits in depressed adolescents. To the best of our knowledge, our study is among the first to use hyperscanning paradigms to examine social deficits in a dynamic context in adolescents with MDD. Interpersonal interaction might be underlined by interpersonal neural synchronization (15). However, many studies only reported the individual neural substrates underlying the social deficits in people with MDD (35), which prevents us from understanding brain function deficits in terms of interpersonal neural interaction. Our current study can fill this gap, demonstrating that hyperscanning paradigms can reveal interpersonal interaction deficits in mental health disorders.

Our study also revealed the INS after the social exclusion was positively correlated with the positive feelings of the other excluded one in the HC group, while this pattern was not observed in the MDD group. In contrast, the changed INS in the MDD group positively correlated with the changed differential pass rate to the partial excluder, while this pattern was not observed in the HC group. These results collectively suggest that the brain-behavioral pattern has somewhat changed in MDD. Such differential brain-behavioral patterns might be informative to differentiate depressed or healthy adolescents. In addition, it also suggests that neuromodulation approaches intended to improve cognitive and affective deficits in individuals with MDD should mind the potentially non-linear relationship between brain signals and behavior.

Our study also has several limitations. Firstly, due to the limited channels of the fNIRS equipment, the signal channels only covered the frontal areas. Future studies could further explore how the neural interaction in other brain regions, including the temporal parietal junction, is involved in social interaction in MDD. Secondly, our experiment setting was restricted to a small group of participants. Future studies could further explore how negative emotions influence social interaction in adolescents with MDD in a setting of a bigger group of participants.

In summary, our study revealed that shared excluded experiences would enhance social interaction preference in adolescents but reduce their INS during social interaction, which indicates adolescents had a reduced willingness to interact with others after social exclusion. Furthermore, the neural interaction in adolescents with MDD was altered to some degree despite no significant behavioral responses being found compared to HC, suggesting the INS might be more sensitive than behavioral responses in probing the social dysfunction in adolescents with MDD. In addition, more effort is necessary to help depressed adolescents deal with psychosocial stressors than healthy adolescents when they might have higher empathic stress under a psychosocial stressor.

## Supporting information

Supplementary methods and results

## Data availability

The datasets generated during the current study are available from the corresponding author upon reasonable request. The codes used for generating the results in this study can be found at Git Hub (https://github.com/hyw5402/Adolescents_SocialExclusionHyperscanning).

## Acknowledgements

The team thanks Kei Kei Lei, Mengze Wang, and Yanyun Lu for their help in recruiting adolescent participants. Thanks to Jingwen Yin, Chi Un Choi, Anqi Gu, Qiwei Guo, Ye Tian, Canpeng Huang, and Songxiang Tang for their help in data collection. Thanks to Dr. Yuhang Long for advice on data analysis. In addition, this work was partly performed at the Super Intelligent Computing Center which SKL-IOTSC, University of Macau supports.

This work was supported by the University of Macau (Nos. MYRG 2020-00067-FHS, MYRG2019-00082-FHS, and MYRG2018-00081-FHS), the Macao Science and Technology Development Fund (No. FDCT0048/2021/AGJ, No. FDCT 0020/2019/AMJ and FDCT 0011/2018/A1).

