## Supplementary methods and results for "Altered Interpersonal Neural Synchronization during Social Interaction After Shared Excluded Experiences in Depressed Adolescents"

**Supplementary materials**

- Supplementary Methods and Materials
- Demographic results
- Supplementary figures
- Trust scale
- Reference

### Supplementary methods and materials

#### Participant exclusion criteria

Participants who had one of the comorbidities or conditions would be removed from the study: substance abuse disorder, severe physical disease, any history of epilepsy before taking the study, or pregnancy. One pair of individuals with MDD dropped out of the study during the task performance, and another five pairs of individuals with MDD were excluded due to incomplete data. Two pairs of healthy controls were excluded due to the incomplete event markers for the imaging recording.

#### Research ethics

The study was reviewed and approved by the Ethics Committee at the University of Macau and the Fifth Affiliated Hospital, Sun Yat-sen University. All participants had provided their written consent prior to their participation in the study.

#### Supplementary paradigm

There were 192 passes in total, consisting of three stages: acceptance stage (two virtual players pass the ball to others without preference, from pass 1 to pass 35), exclusion stage (two virtual players pass the ball between themselves exclusively, from pass 36 to pass 96), and re-acceptance stage (one virtual player, i.e., the partial excluder, passes the ball to others without preference, while the other virtual player, i.e., the full excluder, passes the ball solely to the other virtual player, from pass 97 to pass 192). The re-acceptance stage was further divided into three sessions; each session consisted of 32 passes. The decision-making by the real players was self-paced.

#### Supplementary behavioral statistics of the modified four-player Cyberball game

$$\text{mean pass rate} = \frac{(\text{toss number})_{\text{from1toS}} + (\text{toss number})_{\text{from3toS}}}{(\text{total number})_{\text{from1}} + (\text{total number})_{\text{from3}}}$$

**mean pass rate**

$$\text{mean reciprocal pass rate} = \left(\frac{(\text{toss number})_{\text{from1to3}}}{(\text{total number})_{\text{from3}}} + \frac{(\text{toss number})_{\text{from3to1}}}{(\text{total number})_{\text{from1}}}\right)/2$$

**reciprocal pass rate**

$$\text{diff pass rate} = \frac{\text{abs}\left(\frac{(\text{toss number})_{\text{from1toS}}}{(\text{total number})_{\text{from1}}} - \frac{(\text{toss number})_{\text{from3toS}}}{(\text{total number})_{\text{from3}}}\right)}{\left(\frac{(\text{toss number})_{\text{from1toS}}}{(\text{total number})_{\text{from1}}} + \frac{(\text{toss number})_{\text{from3toS}}}{(\text{total number})_{\text{from3}}}\right)}$$

**differential pass rate**

$$\text{Diff reciprocal pass rate} = \frac{\text{abs}\left(\frac{(\text{toss number})_{\text{from1to3}}}{(\text{total number})_{\text{from3}}} - \frac{(\text{toss number})_{\text{from3to1}}}{(\text{total number})_{\text{from1}}}\right)}{\left(\frac{(\text{toss number})_{\text{from1to3}}}{(\text{total number})_{\text{from3}}} + \frac{(\text{toss number})_{\text{from3to1}}}{(\text{total number})_{\text{from1}}}\right)}$$

**differential reciprocal pass rate**

$$\begin{aligned}\text{skewness} = {} & \text{abs}((\text{pass rate})_{\text{target\_excluded}} - (\text{pass rate})_{\text{target\_part\_excluder}}) \\ & + \text{abs}((\text{pass rate})_{\text{target\_part\_excluder}} - (\text{pass rate})_{\text{target\_full\_excluder}}) \\ & + \text{abs}((\text{pass rate})_{\text{target\_excluded}} - (\text{pass rate})_{\text{target\_full\_excluder}})\end{aligned}$$

**pass rate skewness**

$$\begin{aligned}&\text{meanswitchrate} \\ &= \left(\frac{(\text{No. ofcounts1})_{\text{SwitchFromLastChoice}}}{\text{totalNo. ofchoices1}} + \frac{(\text{No. ofcounts3})_{\text{SwitchFromLastChoice}}}{\text{totalNo. ofchoices3}}\right)/2\end{aligned}$$

**mean switch rate**

The mean pass rate indicates the percentage of ball passes from the excluded participants (player 1 and player 3) to a particular player (S) in totally received balls. The mean reciprocal pass rate indicates the percentage of the ball passes to the other excluded player after receiving the ball from the player. The differential pass rate (diff pass rate) indicates the absolute difference of the pass rate to a particular player (S) between the two excluded participants. The differential reciprocal pass rate (diff reciprocal pass rate) indicates the differential reciprocal pass rate between the two excluded participants. The pass rate skewness indexes the degree of unevenly passing the ball to other players. The mean switch rate indicates the percentage of switching choices in consecutive choices.

### Data acquisition

A BIOPAC fNIRS device (fNIR 2000S) with two sets of 16-channel headbands (channel positions shown in Figure 1. C, formed by 4 light emitters and 10 detectors, with optodes separated by 3 cm) was used to record the hemodynamics of the frontal cortex in two participants simultaneously. It involves light of two wavelengths, including 730 nm and 850 nm. The sampling rate was 10 Hz.

**Data preprocessing**

The original optic density data were converted to changes in concentrations of oxy-hemoglobin (HbO), deoxy-hemoglobin (HbR), and total hemoglobin (HbT) with Modified Beer-Lambert Law (1). A linear detrend method was used to correct the drift noise, followed by motion correction with a wavelet method, and removing the global physiological signals by a principal clustering algorithm (2). A bandpass filtering (0.01-0.8 Hz) was also adopted to exclude the slow fluctuation and the effects caused by physiological signals from heartbeat, respiration, and so on (3).

**Supplementary analysis**

The repeated-measurement ANOVA analysis was conducted to examine whether there were different behavioral response patterns in the sub-sessions of the re-acceptance stage

**Demographic result**

As shown in Table 1, there were no significant differences between the HC and MDD groups in age, gender, and education. In addition, the individuals in the HC and MDD groups had matched working memory capacity. The mean age in the HC group was 17.87 (±1.55) years, while the mean age in the MDD group was 17.21 (±3.06) years. There were 18 male participants and 50 female participants in the HC group, while there were 10 male participants and 58 female participants in the MDD group. The mean educational years in the HC group was 10.90 (±3.02), while that in the MDD group was 11.65 (±1.53).

### Supplementary figures

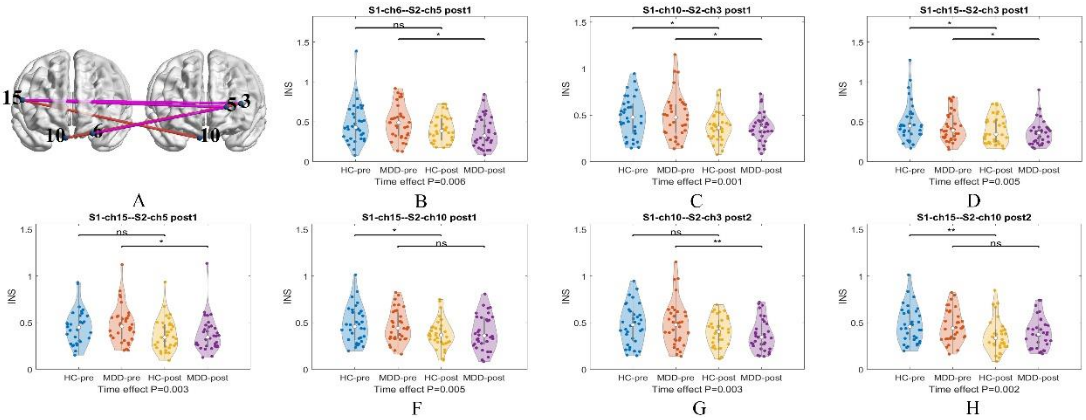


Fig supp 1. The interpersonal neural responses before and after the shared excluded experiences in the separate HC or MDD group.

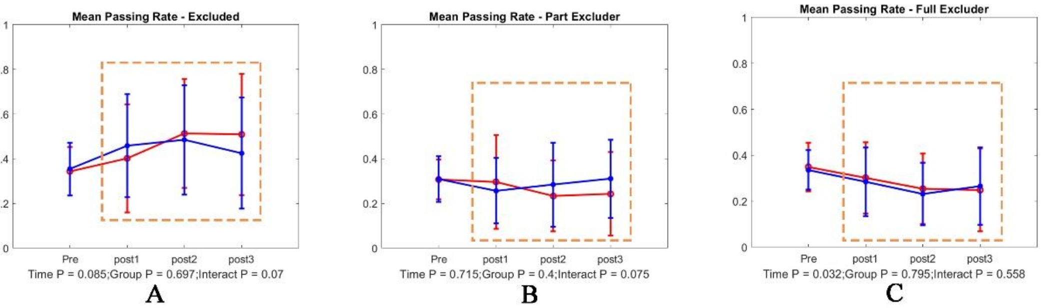


Fig supp 2. Social interaction preference indexed by the mean pass rate showed the trend of time or group-time interaction effects.

**Trust scale**

**Instructions:** Recall the game process you just played and read the following statements. Based on your current feelings, select the appropriate value below to indicate the extent to which you agree or disagree with the statement.

**Strongly Disagree | Partially Disagree | Slightly Disagree | Neutral | Slightly Agree | Partially Agree | Strongly Agree**
**1 | 2 | 3 | 4 | 5 | 6 | 7**

- I am an important member of the game.
- I always think Player1 will pass the ball to me next time.
- I do not feel involved in the game. (R)
- I can gain Player1's trust.
- I look forward to each pass from Player1.
- I do not get much satisfaction from the game. (R)
- Player1 often accompanies me and passes the ball to me.
- I cannot fully trust Player1. (R)
- The number of passes Player1 makes to me and another player is not very different.
- My expectations of receiving passes from Player1 are often disappointed. (R)
- I feel forgotten by Player1. (R)
- If given the chance, I still want to play the game with Player1.
